# Diversification of giant and large eukaryotic dsDNA viruses predated the origin of modern eukaryotes

**DOI:** 10.1101/455816

**Authors:** Julien Guglielmini, Anthony Woo, Mart Krupovic, Patrick Forterre, Morgan Gaia

## Abstract

Giant and large eukaryotic double-stranded DNA viruses from the Nucleo-Cytoplasmic Large DNA Virus (NCLDV) assemblage represent a remarkably diverse and potentially ancient component of the eukaryotic virome. However, their origin(s), evolution and potential roles in the emergence of modern eukaryotes remain a subject of intense debate. Since the characterization of the mimivirus in 2003, many big and giant viruses have been discovered at a steady pace, offering a vast material for evolutionary investigations. In parallel, phylogenetic tools are constantly being improved, offering more rigorous approaches for reconstruction of deep evolutionary history of viruses and their hosts. Here we present robust phylogenetic trees of NCLDVs, based on the 8 most conserved proteins responsible for virion morphogenesis and informational processes. Our results uncover the evolutionary relationships between different NCLDV families and support the existence of two superclades of NCLDVs, each encompassing several families. We present evidence strongly suggesting that the NCLDV core genes, which are involved in both informational processes and virion formation, were acquired vertically from a common ancestor. Among them, the largest subunits of the DNA-dependent RNA polymerase were seemingly transferred from two clades of NCLDVs to proto-eukaryotes, giving rise to two of the three eukaryotic DNA-dependent RNA polymerases. Our results strongly suggest that these transfers and the diversification of NCLDVs predated the emergence of modern eukaryotes, emphasizing the major role of viruses in the evolution of cellular domains.

---

The discovery of giant viruses in the early 21^st^ century has revived the debate on the nature of viruses and their role in evolution^1–13^. The 1µm-long particles of pithoviruses^14^ can be seen under a light microscope and the 2.5Mb-long genomes of pandoraviruses, larger than those of many cellular organisms, encode for more than 2,000 proteins, mostly ORFans^15^. However, these unexpected features notwithstanding, giant viruses are a *bona fide* part of the virosphere, relying on the infected cells for the production of energy and protein synthesis. Phylogenetic and comparative genomics analyses showed that giant viruses together with smaller eukaryotic dsDNA viruses form a supergroup, dubbed the Nucleo-Cytoplasmic Large DNA Viruses (NCLDV)^16,17^. This assemblage encompasses families of large and giant viruses, including *Poxviridae*, *Iridoviridae*, *Ascoviridae*, *Asfarviridae*, *Marseilleviridae*, *Mimiviridae*, and *Phycodnaviridae* as well as several lineages of as yet unclassified viruses, such as pithoviruses, pandoraviruses, molliviruses and faustoviruses^18^. Altogether, the NCLDVs are associated with diverse eukaryotic phyla, from phagotrophic protists to insects and mammals, and some cause devastating diseases, such as smallpox (*Poxviridae*) or swine fever (*Asfarviridae*), or play important ecological roles, such as termination of algal blooms (*Phycodnaviridae*^19^).

The origin and evolution of the NCLDVs remain a subject of controversy. It is still unclear if these viruses form a monophyletic group, if proteins conserved in most NCLDVs had a congruent evolutionary history or if some of them were acquired several times independently from their hosts. Most phylogenetic analyses performed up to now were based on individual proteins or various subsets of conserved proteins^20,21^. These analyses usually recovered the monophyly of various NCLDV families, but often offered contradicting results and the relationships between the families remained debated. For instance, it has been proposed that the giant pandoraviruses are related to members of the *Phycodnaviridae*^22^, but this grouping was not recovered in a recent phylogeny based on their DNA polymerases^23^. According to some studies, the different families of the NCLDVs emerged during the diversification of modern eukaryotes^24^, whereas in other studies, NCLDVs form a monophyletic group branching between Archaea and Eukarya^29^/10/2018 13:51:00. Some authors have even suggested that several families of giant viruses could have originated independently from extinct cellular lineages, possibly even before the last universal common ancestor (LUCA) of Archaea, Bacteria, and Eukarya^11,25^.

With phylogenetic tools being constantly improved and new genomes of large and giant viruses steadily unearthed, we decided to perform an updated and in-depth phylogenetic analysis of the NCLDVs. We mined available genomes for homologous genes, built clusters of orthologous genes, and performed extensive phylogenetic analyses on the 8 most conserved ones, separately and in concatenations. In addition, we have investigated the relationships between NCLDVs and eukaryotes through the phylogeny of the DNA-dependent RNA polymerases (RNAP). Unlike in previous analyses, we included in our study the three eukaryotic RNAP (RNAP I, II, and III) and concatenated their two largest subunits. The robust phylogenies we obtained show that core genes involved in virion morphogenesis as well as genome transcription and replication have co-evolved in the entire NCLDV lineage. Furthermore, our results revealed the existence of two superclades of NCLDVs that diverged after the separation of the archaeal and eukaryotic lineages, but before the emergence of the Last Eukaryotic Common Ancestor (LECA). Surprisingly, our data suggest that eukaryotic RNAP-III is the actual cellular ortholog of the archaeal and bacterial RNAP, while eukaryotic RNAP-II and possibly RNAP-I were transferred between two viral families and proto-eukaryotes. Overall, our results reveal that the diversification of NCLDVs predates the origin of modern eukaryotes: the ancestors of contemporary NCLDVs co-evolved with protoeukaryotes and could have played an important role in the emergence and diversification of modern eukaryotes.

## Results

### Identification of the core genes

Many new NCLDV genomes have been published following the latest comprehensive comparative genomics analyses^21,26^, substantially increasing their known diversity and enriching families that were previously poorly represented. As a result, the list of the most conserved genes among the NCLDVs could have drastically changed since the last estimation, prompting us to re-analyse it. To identify NCLDV orthologs, we designed a pipeline based on Best Bidirectional BLAST Hit combined with manual curation in order to remain as exhaustive as possible while avoiding inclusion of paralogs (see details in Methods section). The sets of conserved proteins classified according to their conservation among NCLDVs are summarized in Supplementary Table 1.

Our results show that only 3 proteins are strictly conserved among the 73 selected NCLDV genomes: family B DNA polymerase (DNApol B), the D5-like primase-helicase (primase hereinafter) and homologs of the Poxvirus Late Transcription Factor VLTF3 (VLTF3-like) (list of genomes in Supplementary Table 2; selection criteria in Methods). Acknowledging various reasons which may preclude detection of homologous genes (e.g., due to high divergence or genuine loss in a taxon), we decided to lower our conservation threshold to include genes found in at least 95% of the genomes. This resulted in the increase of our set of core genes by three: the transcription elongation Factor II-S (TFIIS), the genome packaging ATPase (pATPase), and the major capsid protein (MCP). Notably, no homolog of the MCP has been found in pandoraviruses^15^, whereas pATPases are apparently lacking in Pithovirus^14^, Cedratvirus^27^, and Orpheovirus^28^. Conservation of the NCLDV genes is further discussed in the Supplementary Information.

To this set of six proteins (3 strictly conserved and 3 conserved in 95% of the genomes), we added the two largest RNAP subunits (RNAP-a and -b) despite their notable absence in all genera of the *Phycodnaviridae* family, except for the *Coccolithovirus* genus. Indeed, these two proteins are otherwise highly conserved among the NCLDVs (present in 92% of the genomes) and are the largest universal markers (found in all members of the three cellular domains), which makes them perfectly suited for reconstructing the evolutionary relationships between NCLDVs and cellular organisms. Thus, the set of 8 proteins contains 6 proteins related to informational processes – genomes expression and replication (DNApol B, primase, VLTF3-like, TFIIS, RNAP-a, and RNAP-b) – and 2 proteins involved in virion structure and morphogenesis (pATPase and MCP).

### The core markers share a similar phylogenetic signal

Using a maximum-likelihood (ML) framework, the monophyly of all known NCLDV families, except the *Phycodnaviridae*, was obtained with high support in most of the 8 single-protein phylogenetic trees (Supplementary Figure 1). As often observed in published NCLDV phylogenies^26^, *Ascoviridae* were however nested within the *Iridoviridae* in most trees. The grouping of the *Mimiviridae* with related unclassified viruses with smaller genomes often referred to as the “extended Mimiviridae”^21^ or more recently the “Mesomimivirinae”^29^, was obtained in five out of the 8 trees. We will refer to this grouping as the “Megavirales” putative order (see Supplementary Information).

The *Poxviridae* clade consistently formed a long branch and displayed the most unstable position, branching next to various families (see Supplementary Information). The same was true for *Aureococcus anophagefferens* virus. Thus, to avoid potential artefacts, we decided to remove these taxa from most of our subsequent analyses. Phylogenetic analyses of the resultant dataset resulted in globally congruent trees of individual core proteins (Supplementary Figure 2). Notably, the *Marseilleviridae*, the *Ascoviridae*, the *Iridoviridae*, and a clade grouping Pithovirus sibericum with Cedratvirus A11 and Orpheovirus IHUM-LCC2 (thereafter referred as the Pitho-like viruses), group seemingly together, while the *Phycodnaviridae* (including Pandoraviruses and Mollivirus), *Asfarviridae*, and the “Megavirales” also form a cluster.

In order to verify if the NCLDV informational proteins have indeed co-evolved with proteins involved in virion formation, we first concatenated independently the 4 largest informational proteins (i.e. the DNA and RNA polymerases, and the primase) and next the 2 proteins involved in the formation of virions (the MCP and the pATPase). In both trees (Supplementary Figure 3 and 4), all NCLDV families were monophyletic, except for the *Iridoviridae* which again were split by the *Ascoviridae* in the tree constructed from the concatenation of informational proteins (Supplementary Figure 3). The two phylogenies had similar topologies, with the same clusters of NCLDV families as observed in single-protein trees. Some positions within these clusters might be affected by differences between the two datasets: 2 of the 4 informational proteins are absent in all but one *Phycodnaviridae* genera, while the Pitho-like viruses lack the pATPase gene. The congruence between the two trees still suggests that informational proteins of the NCLDVs have mostly co-evolved with proteins involved in the formation of virions. The 8 core genes hence likely underwent through a similar evolutionary history.

To further confirm that the 8 core proteins have a similar evolutionary history and to detect potential incongruences within the selected proteins that could prevent their global concatenation, we performed a home-made congruence test based on comparative phylogenetic analyses of differential concatenations (see details in Methods; Supplementary Table 3). The topologies of the resulting trees were congruent, with most features systematically present, such as the two clusters of NCLDV families, the presence of groups regularly observed in the ML trees, and the monophyly of families. This test thus did not reveal any major incongruences between the different combinations of core proteins and consequently strongly supports the absence of conflicting signal embedded in a sequence or in a subset of proteins, confirming that the core proteins were likely presents in a common ancestor of NCLDVs and all evolved vertically along their co-evolution with their hosts.

### The evolution of NCLDVs

We concatenated the 8 core proteins together to improve the robustness and resolution of the NCLDV phylogeny. We obtained a ML tree (Supplementary Figure 5) in which the NCLDV families are again clustering into two superclades: the *Marseilleviridae* with the *Ascoviridae,* the Pitho-like viruses’ clade, and the *Iridoviridae* (thereinafter referred as the MAPI superclade), and the *Phycodnaviridae* with the *Asfarviridae* and the “Megavirales” (thereinafter referred as the PAM superclade). All positions in this tree are strongly supported except for the position of the *Asfarviridae* (see Supplementary Information). We further performed Bayesian inferences with the CAT-GTR model, designed to deal with sites and sequences heterogeneity, considering that this could allow a more trustful and accurate reconstruction provided that a satisfactory convergence could be obtained (see Methods). After reaching a good convergence (maxdiff <0.1), we obtained a phylogenetic tree with all nodes at maximum support (Posterior Probabilities = 1), except for two nodes corresponding to minor internal positions within the *Mimiviridae* family. The Bayesian tree was almost identical to the ML tree, except that *Phycodnaviridae* are now sister group to a clade clustering *Asfarviridae* and “Megavirales” (Fig 1). This topology was also confirmed using a supertree approach (Supplementary Figure 6; details in Methods and Supplementary Information).

**Fig 1.**
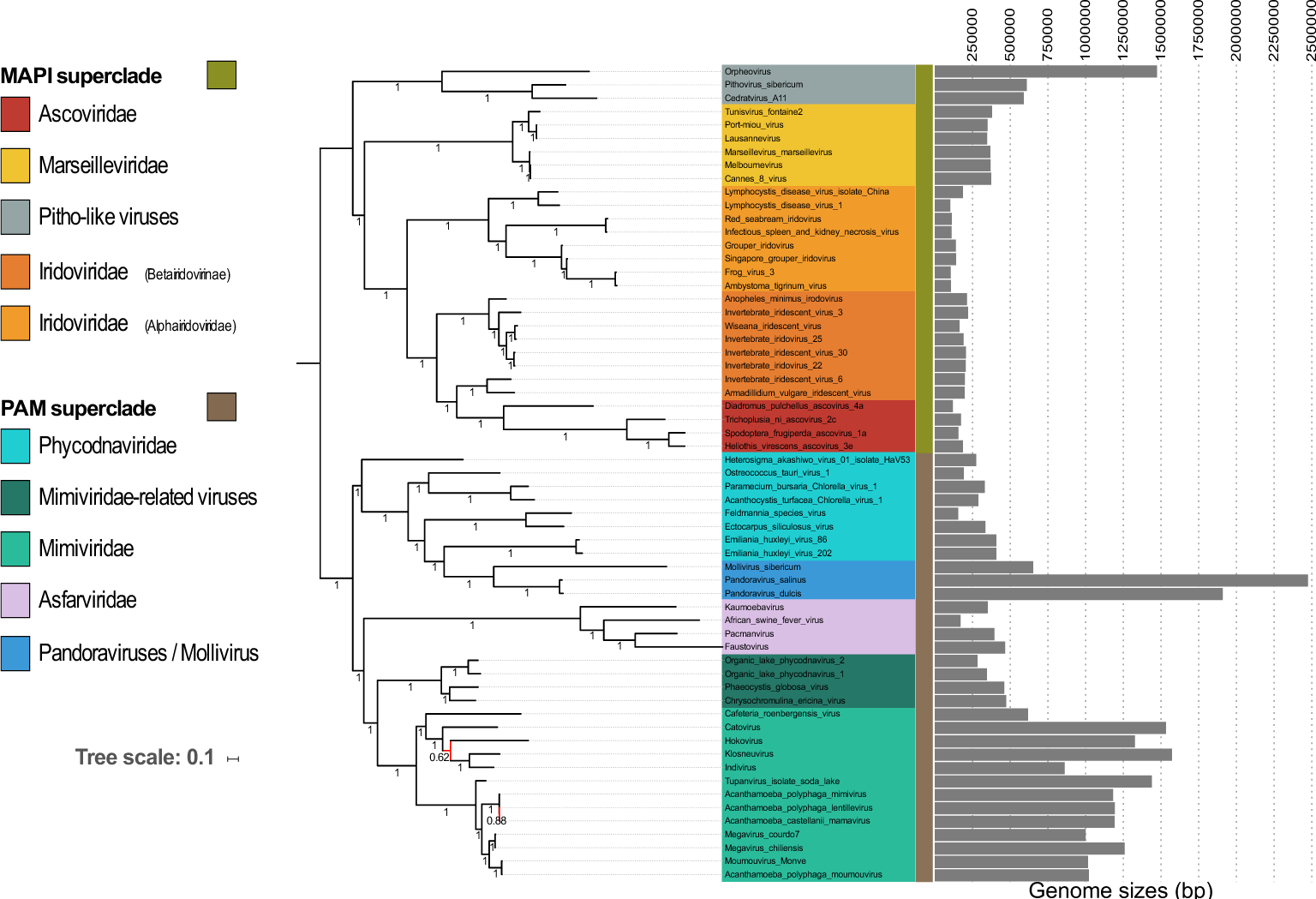
Phylogenetic tree of the NCLDVs. Bayesian inference (CAT-GTR model) of the concatenated 8 core proteins from the NCLDVs after removal of *Poxviridae* and *Aureococcus anophagefferens* virus. Genome sizes (in bp) are represented next to each virus name. The scale-bar indicates the average number of substitutions per site. The values at branches represent Bayesian posterior probabilities. Nodes without maximum support are indicated in red.

This tree confidently positions recently identified viruses. The *Mimiviridae* hence include Klosneuvirus, Indivirus, Catovirus, Hokovirus^30^, and Tupanvirus^31^, and are associated with related viruses within the putative “Megavirales” order. The still unclassified Pitho-like viruses, which herein consists of Pithovirus sibericum, Cedratvirus A11, and Orpheovirus IHUM-LCC2, seem to represent a new separate family whose position within the putative MAPI superclade remains to be investigated to further extent considering their still low representation. Faustovirus^32,33^, Pacmanvirus^34^, and Kaumoebavirus^35^, form a well-supported clade with the African swine fever virus (ASFV-1) of the *Asfarviridae*, as previously suggested^36^. The *Phycodnaviridae* encompass pandoraviruses and Mollivirus sibericum. The monophyly of this family however remains a matter of debate as it is not observed in half of the single-protein trees and has low support in the ML tree based on the concatenated structural proteins. This is possibly due to the very large diversity of the viruses within this family. Altogether, our in-depth phylogenetic analyses nonetheless strongly support the existence of the two major superclades, the MAPI and the PAM.

The evolution and origin of NCLDVs is regularly debated, most notably in term of their connections to other viruses^18^. Interestingly, homologs of the MCP and pATPase can be found in viruses from various families belonging to the PRD1-Adenovirus lineage. This lineage was initially proposed based on the structural conservation of the major capsid proteins as well as shared principles of virion assembly and genome packaging^37–39^. The closest outgroup to NCLDVs in this lineage could be Polintoviruses^40,41^. When using Polintoviruses as an outgroup (see Methods), the ML tree of the MCP-pATPase concatenation is split between the MAPI and PAM putative superclades, suggesting that these two clusters indeed form monophyletic assemblages (Fig 2). Notably, the MCP-pATPase tree remains almost identical to the one obtained with the NCLDVs alone (the only difference being the position of the *Phycodnaviridae*), and the number of positions was not dramatically reduced (601 positions with Polintoviruses versus 625 positions without). This indicates that the split between the MAPI and PAM superclades was probably the earliest event in the evolution of known modern NCLDVs from their common ancestor.

**Fig 2.**
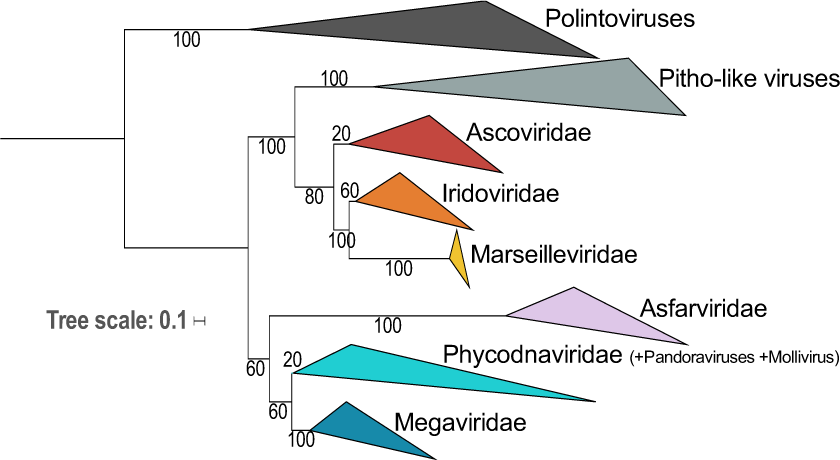
Relationships between Polintoviruses and NCLDVs. Maximum likelihood (ML) phylogenetic tree of the concatenated structural proteins from Polintoviruses and NCLDVs after removal of *Poxviridae* and *Aureococcus anophagefferens* virus. The scale-bar indicates the average number of substitutions per site. The values at branches represent support calculated by nonparametric bootstrap.

### The relationship between NCLDVs and the three cellular domains

The RNA and DNA polymerases of NCLDV have homologues in the three domains of life (Archaea, Bacteria and Eukarya), making it *a priori* possible to investigate their evolutionary relationships with cellular organisms. However, the family B DNA polymerase, often used to tentatively affiliate new NCLDV genomes to known taxa^42^, cannot be used for this task since they are absent from most Bacteria and their phylogenetic analyses produce complex scenarios with the two major subgroups of archaeal DNA polymerases intermingled with the four types of eukaryotic family B DNA polymerases (α, δ, ε, ζ)^43^. In contrast, phylogeny of the two largest RNAP subunits, which are also the largest universal markers, recovered the monophyly of the three cellular domains^44^. Thus, RNAPs are good candidates to study the relationships between the cellular domains and NCLDVs.

Most phylogenetic analyses of RNAPs performed until now included only the eukaryotic RNA polymerase II (RNAP-II), which is the most studied and usually considered as the most similar to the archaeal RNAPs^45^. Here, we decided to include all three eukaryotic RNAPs (RNAP-I, RNAP-II and RNAP-III) (we used a normalized nomenclature, see Supplementary Information). Importantly, these three multi-subunit RNAPs are present in all eukaryotes, indicating that they were already all present in the Last Eukaryotic Common Ancestor (LECA). Their inclusion in our dataset thus should both reduce the length of the eukaryotic branch and provide three universal eukaryotic phylogenies, thus three positions for LECA in the cellular/NCLDV RNAP tree.

We have previously obtained a robust phylogenetic RNAP tree with a concatenation of the two largest RNAP subunits (in ML and Bayesian frameworks), in which the three domains are monophyletic, with Eukaryotes and Archaea being sister groups (the so-called Woese’s tree). We obtained this result using a balanced dataset (same number of species for each of the three domains) and avoiding known fast-evolving species to prevent long branch attraction artefacts^44,46^29/10/2018 13:51:00. Since our initial dataset included only RNAP-II as the eukaryotic representative, we added the eukaryotic RNAP-I and RNAP-III (list of selected taxa in Supplementary Table 4). Interestingly, Archaea and Eukarya again form two monophyletic sister groups in our new concatenated RNAP subunits tree, despite the drastic reduction of the eukaryotic branch length (Supplementary Figure 7). Remarkably, RNAP-I was not attracted by Bacteria despite its very long branch. These observations suggest that the three-domain topology of the RNAP tree did not result from the attraction of eukaryotes by the long bacterial branch. Interestingly, the three eukaryotic RNAPs displayed globally congruent phylogenies, corroborating their presence in LECA.

We included the sequences of NCLDVs into this new dataset (except for *Poxviridae* and *Aureococcus anophagefferens* virus) in order to investigate the timeline of NCLDVs diversification in the context of cellular evolution. The ML phylogenetic analysis of concatenated RNAP subunits yielded the three-domain topology (Supplementary Figure 8) in which NCLDVs branch after the divergence of the archaeal and eukaryotic lineages. We then removed Bacteria from our subsequent analyses in order to increase the resolution (single-protein trees in Fig 3 and in Supplementary Figure 9; concatenation in Supplementary Figure 10). The trees were highly similar after selecting the Archaea as the outgroup, and supports for several nodes indeed became stronger. Since each of the cellular clades (the Archaea and the three eukaryotic homologs) was well represented and systematically monophyletic, we decided to use the cellular sequences as constraints during the alignment process (each of the 4 clades of cellular sequences corresponding to an independent constraint; see details in Methods), allowing us to check if this could improve the resolution by limiting mis-alignments from small insertions or deletions in the viral sequences. The resulting concatenation of the two subunits switched from 1,683 positions to 1,595, and the highly supported reconstructed tree obtained in ML framework (LG+C60 model) (Fig 4) was strictly identical to the one without any constraint. The most significant feature of the viral/cellular RNAP tree is that LECA, despite being a single timepoint in the history of eukaryotes, is represented three times among the diversity of NCLDVs, indicating that NCLDVs predated LECA. This reveals that the diversification of NCLDVs itself predated that of modern eukaryotes, and consequently, different NCLDV families or superclades were already infecting proto-eukaryotes.

**Fig 3.**
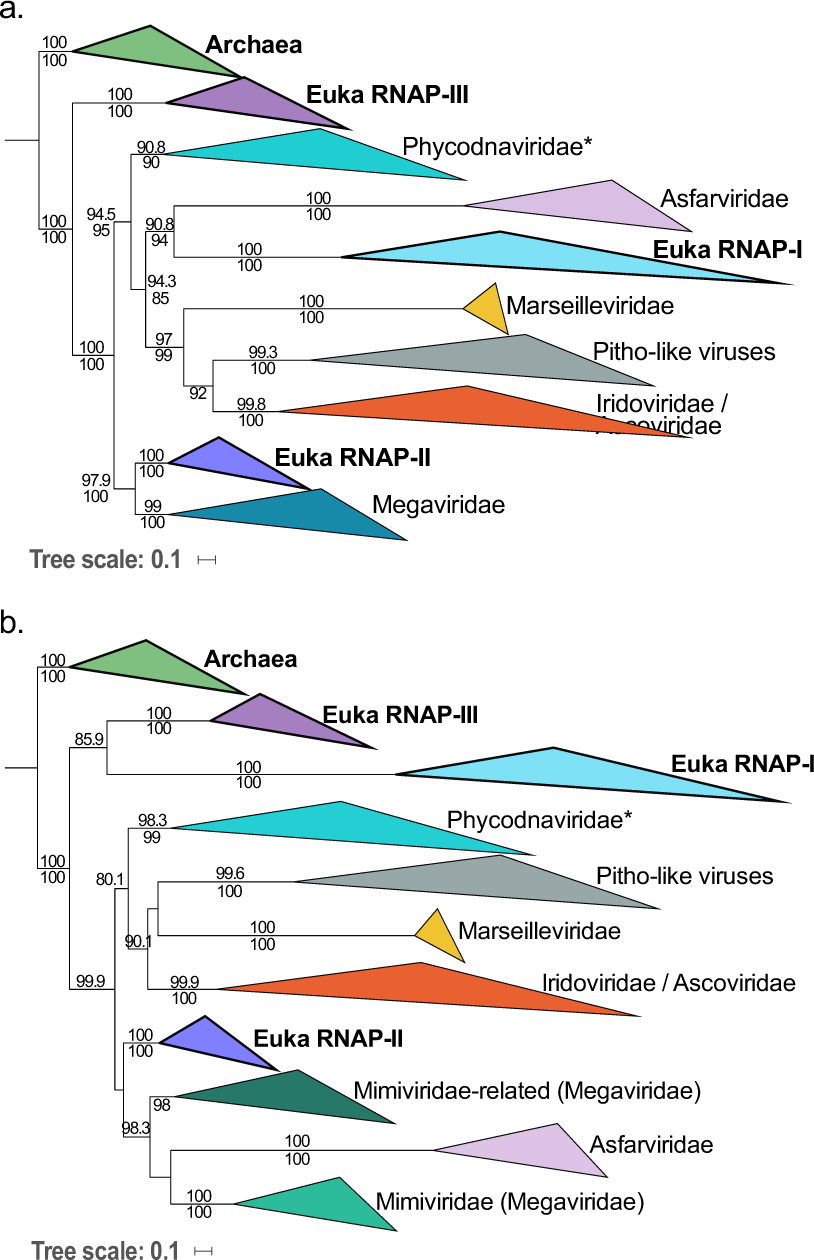
Maximum likelihood (ML) single-protein trees of the two largest RNA polymerase subunits from Archaea, Eukaryotes, and NCLDVs. ML phylogenetic trees of the RNAP-a (**a**) and RNAP-b (**b**) subunits, with Archaea used as the outgroup. The scale-bars indicate the average number of substitutions per site. Values on top and below branches represent support calculated by SH-like approximate likelihood ratio test (aLRT; 1,000 replicates) and ultrafast bootstrap approximation (UFBoot; 1,000 replicates), respectively. Only values superior to 80 are shown.

Surprisingly, in the tree based on concatenated RNAP subunits, the eukaryotic RNAP-III appears to be the closest to the archaeal outgroup after addition of viral sequences with strong supports, suggesting that it could be the actual ortholog of the archaeal enzyme (Fig 4). A major feature of this tree is that NCLDVs do not form a monophyletic group, but three monophyletic subgroups well separated from the three eukaryotic RNAPs, instead of emerging from within eukaryotic diversity. In order to test this result, we performed an Approximately Unbiased (AU) tree topology test and compare this tree to two others constraining either the monophyly of NCLDVs or cellular organisms (see Methods). The AU test rejected these two alternative trees with p-values <1e-3. Remarkably, the relative positions of the NCLDV families and superclades in the RNAP tree are completely congruent with the NCLDV topology in the Bayesian tree previously obtained with the 8 core proteins (Fig 1) and highly similar to the tree obtained using the concatenation from which the two RNAP subunits were omitted during the congruence test (Supplementary Table 3; Supplementary Figure 11). In particular, we recovered the monophyly of the MAPI superclade, and its internal phylogeny is highly similar to that obtained previously (the positions of *Marseilleviridae* and Pitho-like viruses are flipped).

**Fig 4.**
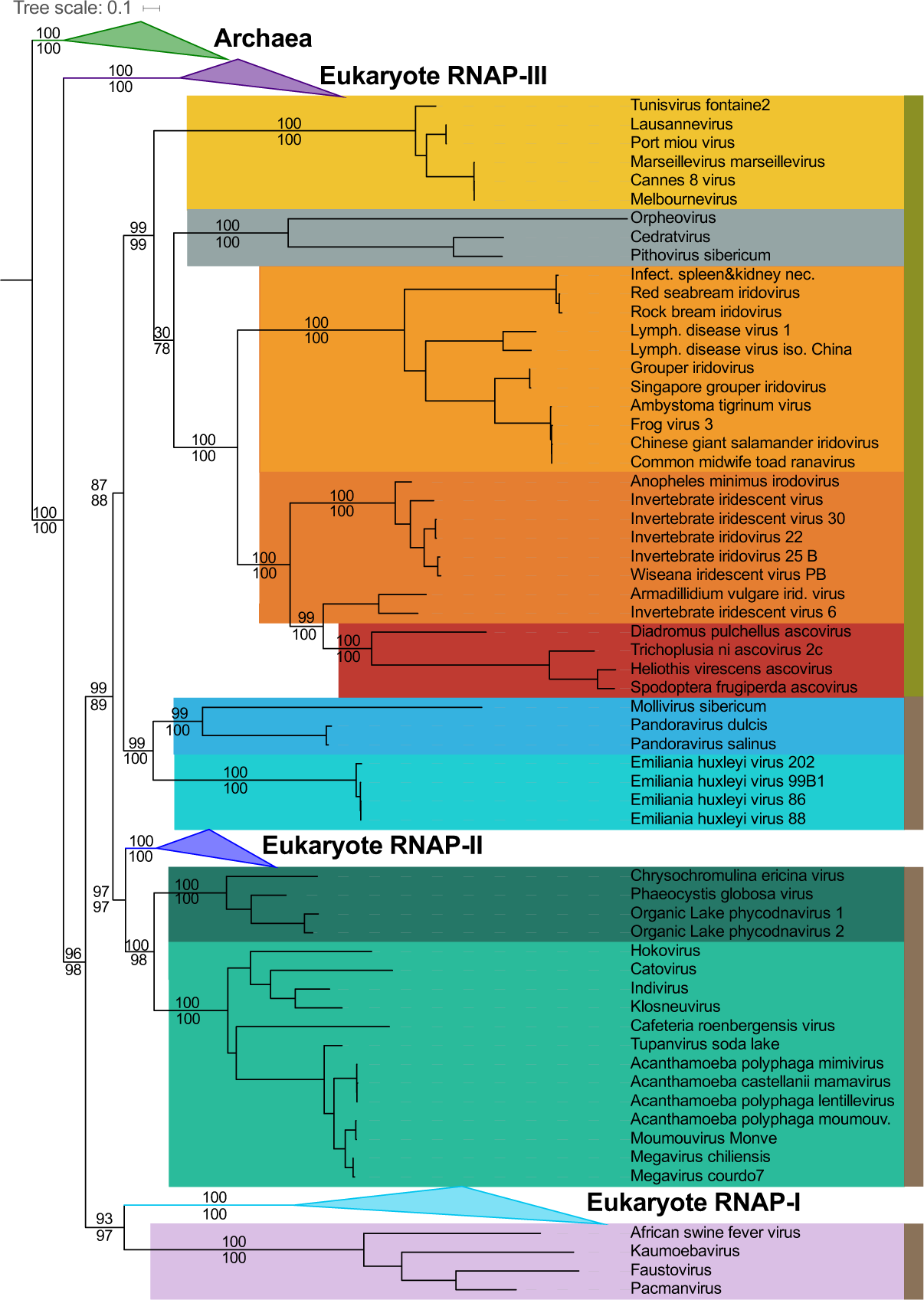
Maximum likelihood (ML) phylogenetic tree of the concatenated two largest RNAP subunits from Archaea, Eukaryotes, and NCLDVs. ML phylogenetic tree of the concatenation of the two largest RNAP subunits, with Archaea used as the outgroup. Among the PAM superclade (light brown), “Megavirales”, *Asfarviridae*, and *Phycodnaviridae* are indicated in light/dark green, pink, and light/dark blue, respectively. Among the MAPI superclade (olive green), the *Marseilleviridae*, Pitho-like viruses, *Iridoviridae*, and *Ascoviridae* are indicated in dark yellow, grey, light/dark orange, and red, respectively. The scale-bar indicates the average number of substitutions per site. Values on top and below branches represent support calculated by SH-like approximate likelihood ratio test (aLRT; 1,000 replicates) and ultrafast bootstrap approximation (UFBoot; 1,000 replicates), respectively.

Four clades of the NCLDVs are distinguishable in this viral-cellular RNAP tree, corresponding to the monophyletic MAPI superclade, the *Phycodnavirida*e, the “Megavirales” and the *Asfarviridae*. The PAM superclade is indeed not monophyletic in the RNAP tree because eukaryotic RNAP-I and -II are branching within it. The relative positions of the three PAM families compared to each other are still matching the NCLDV tree topology obtained with the 8 core proteins in the Bayesian framework (Fig 1), but in the viral/cellular RNAP tree, the eukaryotic RNAP-II is sister group to the “Megavirales” whereas the eukaryotic RNAP-I is sister group to *Asfarviridae*. In order to assess the robustness of these groupings, and notably of the *Asfarviridae* and RNAP-I that both display long branches, we reconstructed a consensus bootstrap tree of the concatenated RNAP subunits. In parallel, we also performed a phylogenetic analysis based on reconstructed ancestral sequences to replace the three eukaryotic RNAP clades (see Methods). Both methods supported the relationships between the “Megavirales” and the eukaryotic RNAP-II as well as between the *Asfarviridae* and the eukaryotic RNAP-I, suggesting that they reflect a genuine evolutionary signal (Supplementary Figure 12). Worth-noting, the position of the *Asfarviridae* differs in the two single-protein subunit trees: they are sister group to the RNAP-I in the individual *a* subunit tree (Fig 3a), as in the tree based on concatenated RNAP subunits (Fig 4), whereas they branch within the “Megavirales” in the *b* subunit tree (Fig 3b). This suggests that two transfers might have occurred between proto-eukaryotes and ancestors of the *Asfarviridae* and could explain the long branch of the *Asfarviridae* in the RNAP trees.

Considering the branching of NCLDVs after the eukaryotic RNAP-III, it seems that they have originally obtained their RNAP from proto-eukaryotes after their divergence from the archaeal lineage. The unexpected positions of RNAP-I and -II within NCLDVs could suggest that these two eukaryotic RNAPs were either recruited from NCLDVs or transferred to the ancestors of the *Asfarviridae* family and “Megavirales” order. The latter hypothesis seems unlikely because replacements of the two largest core genes of two major NCLDV families by their cellular counterparts would have likely resulted in substantial alterations in the NCLDV topologies obtained during the congruence test. This was not the case, and notably, the tree produced without RNAP genes during this test (Supplementary Figure 12) was highly similar with the 8-core-proteins tree (Fig 1), and with the trees from the concatenated RNAP genes only, with (Fig 4) or without cells (Supplementary Figure 13). The only difference is the position of *Phycodnaviridae*, which are sister group to “Megavirales” in the absence of RNAP genes. This is remarkable since the RNAP proteins represent nearly half of the total positions in the global concatenation. These data strongly suggest that the transfers of the RNAP-encoding genes were directed from viruses to cells, after the diversification of these RNAPs within NCLDVs. Based on this observation, we postulate a possible scenario depicted in Fig 5. In this hypothesis, the ancestral eukaryotic RNAP (at least the two largest subunits), more similar to RNAP-III, was first transferred to the ancestor of NCLDVs. After the divergence between the MAPI and the PAM superclades, this viral RNAP diverged in the common ancestor of “Megavirales” and *Asfarviridae*, and was transferred to proto-eukaryotes, later to become the RNAP-II. Separately, a duplication of the ancestral RNAP-III in proto-eukaryotes occurred, before the largest subunit of this newly formed RNAP was replaced by that of *Asfarviridae*: this new complexe, partly viral and partly cellular from duplication, resulted in the RNAP-I.

**Fig 5.**
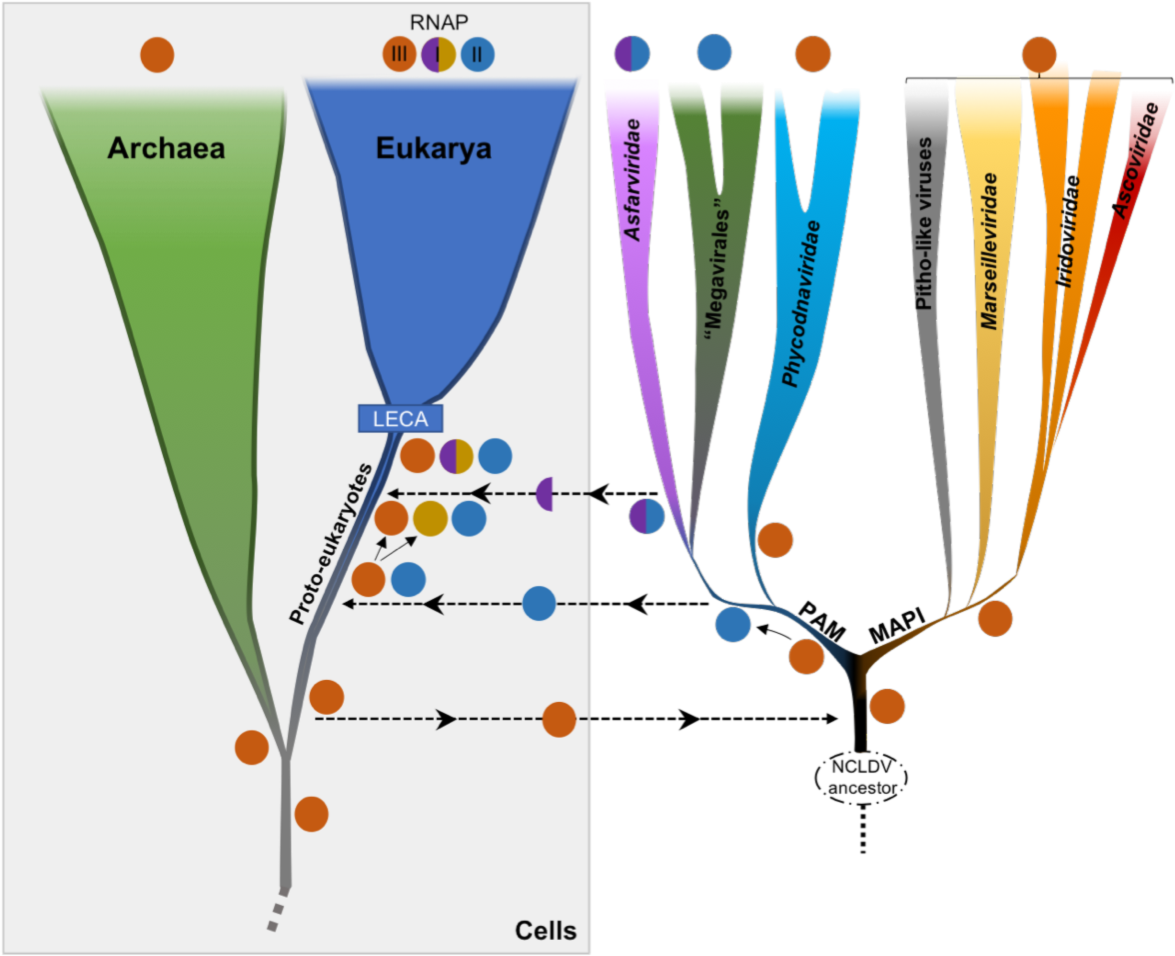
Schematic representation of a putative scenario for the transfers of RNAP between cells and NCLDVs. An ancestral RNAP that later gave rise to the eukaryotic RNAP-III, actual ortholog of the archaeal RNAP, was transferred (at least the two largest subunits) from proto-eukaryotes to the ancestor of modern NCLDVs. A significantly divergent RNAP was later on transferred from the common ancestor of *Asfarviridae* and “Megavirales” to proto-eukaryotes. A new eukaryotic RNAP also emerged from a duplication event from the RNAP-III, before its largest subunit was replaced by that of *Asfarviridae*. These events occurred before LECA, the Last Eukaryotic Common Ancestor, that marked the emergence of modern eukaryotes.

## Discussion

From our investigation of the NCLDV genomes, including those of most recently identified giant and large dsDNA viruses, we could reconstruct a robust phylogenetic tree of this group likely to represent their vertical evolutionary history. Our results provide a solid framework for proposed and sometimes debated positions of different NCLDV families. Notably, Pithovirus and related viruses form a separate, yet to be named family most closely related to the *Marseilleviridae*. Pandoraviruses and Mollivirus branch within the *Phycodnaviridae*, as a sister group to *Coccolithovirus* genus, confirming the results of Yutin and Koonin^22^. Our results reveal two robust monophyletic superclades, the MAPI and the PAM, each of which includes several virus families and a number of unclassified viruses. These results call for reassessment of the taxonomy of large and giant dsDNA viruses included in the NCLDV assemblage. In particular, the expansion of the *Mimiviridae* family and discovery of associated but more distantly related viruses suggests that a family-level taxon might not be adequate to encompass this diversity. Consequently, the *Mimiviridae* and the related algal viruses as well as viruses discovered by metagenomics might have to be unified into a new order, the “Megavirales”. Furthermore, the *Asfarviridae* clade, in addition to ASFV-1, includes the Faustovirus^32,33^, Kaumoebavirus^35^ and Pacmanvirus^34^, which have been suggested to represent separate families^35^. Thus, an order-level taxon would be needed for classification of these viruses. Similarly, in the MAPI superclade, the placement of the pandoraviruses and the mollivirus within the *Phycodnaviridae* indicates that this family might not be monophyletic and should be revised. *Ascoviridae* regularly branch within *Iridoviridae*, advocating for a reconsideration of these two families. The elusive position of the *Poxviridae*, which were removed from most of our analyses, and their actual association to NCLDVs remain to be investigated.

The monophyly of NCLDVs is not recovered in the cellular/NCLDV RNAP tree: NCLDVs do not form a fourth domain of life, as proposed by some^20^, nor nest among eukaryotes^24^. While some genes in the NCLDV genomes might have been recruited from different sources, notably their modern hosts and bacteria, we have shown that a congruent vertical evolutionary history of NCLDVs is traceable and sound. The 8 selected core genes selected indeed shared a similar vertical evolution, and were inherited from a common ancestor, which was likely smaller, as hypothesized before^47^, and specifically related to polintoviruses^12^. Notably, these core genes are involved in both genome replication and virion formation, key features of viruses, supporting their evolution from a viral ancestor. The division into the two superclades that our results confidently describe seems to have been the most basal event in the evolutionary history from this ancestor toward modern NCLDVs. The MAPI superclade gave rise to *Marseilleviridae*, *Ascoviridae*, Pitho-like viruses, and *Iridoviridae*. The second superclade, PAM, comprises the *Phycodnaviridae*, the *Asfarviridae*, and the “Megavirales”. Interestingly, giant viruses do not cluster together in the NCLDV trees. Most of them are present in the PAM superclade, but in two separate families (*Mimiviridae* and *Phycodnaviridae*), whereas Orpheovirus is present in the MAPI superclade (Fig 1). The scattered distribution of giant viruses within the diversity of NCLDVs strongly opposes a giant – viral or cellular – ancestor scenario as proposed previously^11,25^. By contrast, it suggests that along the evolution of NCLDVs massive increases in genome size have occurred several times independently in different virus groups, potentially through successive steps of reduction and expansion of their genomes^48,49^.

Our analyses of the two largest subunits of the RNAP, including the three eukaryotic polymerases, revealed that the genuine ortholog of the archaeal and bacterial RNAP might actually be the eukaryotic RNAP-III. In agreement with this unexpected result, homologs of the eukaryotic RNAP-III specific subunit RPC34 are present in most archaeal lineages^50,51^. Importantly, the inclusion in our analyses of the three eukaryotic polymerases, which emerged and were fixed in the LECA before the emergence of modern eukaryotes, provided a relative timeframe for the NCLDVs’ origin and diversification. Our RNAP trees, by positioning the three monophyletic eukaryotic homologs, representing LECA, within the diversity of NCLDV families strongly imply that the evolution of NCLDVs toward the MAPI and PAM superclades and subsequent emergence of the constituent families predated the evolutionary bottleneck that marked the emergence of modern eukaryotes. Several authors have suggested that NCLDVs have played a central role in the origin of eukaryotes^7,9,52–54^. Our results indeed suggest that modern eukaryotes obtained two of their three RNAP, RNAP-I and RNAP-II from NCLDVs. Preliminary studies also suggested that eukaryotes obtained their major type II DNA topoisomerases from NCLDVs^55^. It will be interesting to test these enzymes as alternative outgroups to root the eukaryotic tree. Our results indicate that further digging into the diversity and molecular biology of NCLDV will probably have a major impact on our understanding of the origin and early evolution of eukaryotes.

## Methods

### Datasets

We initially collected a total of 96 NCLDV genomes from public databases (Supplementary Table 2) that we used to build their core genome (see below). This dataset comprises 17 Mimiviridae, 6 Marseilleviruses, 30 Iridoviridae, 4 Ascoviridae, 14 Poxviridae, 4 Asfarviridae,15 Phycodnaviridae, 3 unclassified viruses (referred to as Pitho-like viruses), 2 Pandoraviruses,1 Mollivirus.

Preliminary phylogenetic analyses showed high redundancy within some groups already comprising many members compared to others. We thus decided to remove some genomes in order to obtain a more balanced sampling (Supplementary Table 2): 14 *Iridoviridae*, 2 *Phycodnaviridae* and 4 *Mimiviridae*. These analyses also revealed that the *Poxviridae* on the one hand, and a single virus (*Aureococcus anophagefferens virus*) on the other hand, always produce long branches and tend to change position in the tree depending on the considered proteins or concatenation of proteins. We thus decided to remove these viruses (14 *Poxviridae* and *Aureococcus anophagefferens virus*) from subsequent analyses, leading to the dataset of 61 genomes used in the phylogenetic analyses.

Ten polintoviruses sequences were collected from the Repbase collection^56^ (http://www.girinst.org/Repbase_Update.html): Polinton-1_HM, Polinton-3_TC, Polinton-5_NV, Polinton-2_NV, Polinton-1_DY, Polinton-1_TC, Polinton-1_SP, Polinton-2_SP, Polinton-2_DR, Polinton-1_DR.

The cellular taxa included in some analyses were selected based on previous works performed by some of us^44^. The list of selected taxa is presented as Supplementary Table 4.

### Core genome building

Because of the high divergence level of NCLDV genomes, we were not able to directly identify genes shared among all of them. This is why we first started from two subsets of NCLDVs, both being coherent enough and comprising enough members. Those two subsets were the viruses annotated as *Mimiviridae* on the one hand and *Marseilleviridae* on the other hand.

For each subset of genomes, we proceeded as follow. We defined groups of orthologous genes by blasting one proteome against all the others. We only considered hits that had an E-value less than 1e^-10^. We then identified pairwise reciprocal best hits with at least 20% similarity, and at least 40% of alignment coverage. We finally identified the union of all the sets of orthologs and retained those present in more than half of the members of the subset.

The result was two sets of orthologs, one for each subset of NCLDVs genomes. We compared these two sets by identifying the matching proteins using BLAST and HMM profiles and obtained orthologs found in both *Mimiviridae* and *Marseilleviridae*. Using the aforementioned BLAST criteria, we checked for the presence of these orthologs in other NCLDVs proteomes. When a protein was missing, we checked the presence of a corresponding gene using TBLASTN to account for incomplete annotations of the genomes, and also used HMM profiles to account for high sequence divergence. This whole process resulted in a set of putative orthologous proteins found in all NCLDV families.

In order to detect errors, typically different proteins assigned to the same group, we used HMMer^57^ to find a matching HMM profile in the PFAM database (http://pfam.xfam.org/) for each group and discarded those significantly matching more than one PFAM profile (after checking that these profiles were not from the same protein family). We finally aligned the remaining orthologs and visually inspected the alignments as a last control.

We obtained a list of orthologs that we ordered according to their presence in NCLDV genomes to define different categories of core proteins.

### Phylogenetic analyses

#### Alignments

All alignments were performed using MAFFT v7.397 and the E-INS-i algorithm^58^, which is designed to align sequences that are susceptible to contain large insertions. For one RNA polymerase analysis (see manuscript), constraints in the alignments were used with the seed option: independent alignments of each cellular clade (Archaea and the three eukaryotic RNA polymerases) performed separately were used as constraints for the global alignment. For the viral phylogenies, we trimmed each alignment of the positions containing more than 20% of gaps using our own scripts. For the RNA polymerase phylogenies with cellular sequences, the alignments were trimmed with BMGE (with the -m BLOSUM30 and -b 1 options)^59^.

#### Maximum likelihood phylogenies

Single-protein and concatenated protein phylogenies were conducted within the Maximum Likelihood (ML) framework using IQ-TREE v1.6.3^60^. We first performed a model test with the Bayesian Information Criterion (BIC) by including protein mixture models^61^. For mixture model analyses, we used the PMSF models^62^. The support values were either computed from 100 bootstrap replicates in the case of nonparametric bootstrap, or from 1,000 replicates for SH-like approximation likelihood ratio test (aLRT)^63^ and ultrafast bootstrap approximation (UFBoot)^64^.

#### Congruence analysis

To detect potential incongruences within the signal carried by core proteins (after removal of Poxviridae and Aureococcus anophagefferens virus) that could prevent their global concatenation, we performed comparative phylogenetic analyses of every possible combinations of 6 out of 8 core proteins through ML framework (see ML method aforementioned). The 36 ML trees generated were carefully analyzed for reference features estimated from the Bayesian phylogenetic tree (Fig 1), as well as from most phylogenetic trees obtained throughout this study. The presence or absence of these features were counted, and accordingly each feature was scored for its observed frequency among the trees, as well as each tree was scored according to the number of observed reference features (Supplementary Table 3).

#### Supermatrix analysis

We obtained a supermatrix by concatenating the 8 amino acid alignments of the core genes. Supermatrices containing more characters, we computed ML trees with the aforementioned method and performed Bayesian analyses using phyloBayes MPI v1.5a^65^ and the CAT-GTR model^66^. Four independent chains were run until at least two reached convergence with a maximum difference value <0.1. The tree presented in Fig 1 was obtained from the convergence (maxdiff value: 0.097) of two chains of 3,426 and 3,276 generations. The first 25% of trees were removed as burn-in. The consensus tree was obtained by selecting one out of every two trees. In order to account for composition bias, we also applied two different character recodings, using 4 bins according to two different binnings: the adaptation of the 6 Dayhoff groups^67^ to 4 bins proposed by Lartillot in phyloBayes manual, and the one proposed by Susko and Rogers^68^. For these analyses, a GTR+Γ_4_+I model was used.

#### Supertree analysis

Horizontal gene transfers can deeply impact tree reconstruction when using alignment-based methods. Supertree methods aim at reconciliating sets of phylogenetic trees, typically gene/protein trees, into an organismal tree even when such evolutionary phenomena occur. Among the different proposed criteria for supertree methods, the subtree prune-and-regraft (SPR) distance has proven to lead to more accurate tree reconstructions^69^. We used the software SPR Supertree v1.2.1^69^ from the 8 single protein phylogenies we previously inferred, after collapsing the clades for which the support was less than 95%.

#### Ancestral sequence reconstruction

In order to try to reduce the risk of long branch attraction, we replaced, in the RNAP tree, the eukaryotic clades by their ancestral sequences. These sequences were inferred using IQ-TREE. We selected sites with a posterior probability greater than 0.7 and replace the other sites by gaps.

#### Topology test

IQ-TREE v1.6.3 was used to perform Approximately Unbiased (AU) tree topology tests^70^ for comparing the tree obtained with the concatenated RNAP genes (Fig 4) with two other ones we built using the same methodology but constraining i) the monophyly of the NCLDVs and ii) the monophyly of the cellular organisms. The AU tests rejected these two new trees with p-values <1e-3.

#### Visualization

The phylogenetic trees were visualized with FigTree v1.4.3 (http://tree.bio.ed.ac.uk/software/figtree/) and iTOL^71^.

